# A reference genome assembly of the alpine forage grass *Elymus nutans*

**DOI:** 10.1101/2025.03.29.645954

**Authors:** Dan Chang, Shangang Jia, Ming Sun, Tao Huang, Huanhuan Lu, Jiajun Yan, Changbing Zhang, Minghong You, Jianbo Zhang, Lijun Yan, Wenlong Gou, Xiong Lei, Xiaofei Ji, Yingzhu Li, Decai Mao, Qi Wu, Ping Li, Hongkun Zheng, Xiao Ma, Xuebin Yan, Quanlan Liu, Xiaofan He, Wengang Xie, Daxu Li, Shiqie Bai

**Affiliations:** Sichuan Academy of Grassland Sciences, Chengdu, China; School of Life Science and Engineering, Southwest University of Science and Technology, Mianyang, China; College of Pastoral Agriculture Science and Technology, Lanzhou University, Lanzhou, China; College of Grassland Science and Technology, China Agricultural University, Beijing, China; Biomarker Technologies Corporation, Beijing, China; College of Animal Science, Guizhou University, Guiyang, China; College of Grassland Science and Technology, Sichuan Agricultural University, Chengdu, China; College of Animal Science and Technology, Yangzhou University, Yangzhou, China; College of Biological Engineering, Qingdao University of Science & Technology, Qingdao, China; College of Grassland Science, Gansu Agricultural University, Lanzhou, China

## Abstract

*Elymus nutans* Griseb. (Poaceae: Triticeae, 2n=6x=42) is a dominant perennial plant species in the Qinghai-Tibetan Plateau in China, which is an important forage resource in high-altitude and cold regions and is the most popular species used for high yield artificial grassland planting because of its rich nutritional value and good palatability for herbivorous ruminant animals. In this study, using advanced sequencing technology, we generated an allohexaploid reference genome for *E. nutans*, representing the three sets of chromosomes (subgenomes St, Y, and H). This is the first study to our knowledge to confirm the origin of the Y subgenome, which shares a close relationship with the V subgenome. We predict that *E. nutans* arose via two-step hybridization and that its speciation occurred ∼3.16 MYA. The whole genome in this study will update the Triticeae family genomics information and provide theoretical basis for the evolutionary of its related speicies.

## Introduction

*Elymus nutans* Griseb. (Poaceae: Triticeae, 2n=6x=42) is a dominant perennial plant species (Fig. 1A) in the Qinghai-Tibetan Plateau in China (Chen et al., 2009; Liu et al., 2022), where it serves as an important forage grass with high yields, high nutritional value, and good palatability for herbivorous ruminant animals. At present, the research of E.nutans were mainly focused on domestication and breeding, seed production, artificial cultivation technologies, genetic variation, and both biotic and abiotic stress resistance. Due to the lack of in-depth molecular research as the foundation, some important molecular mechanisms for it have not been reported, which caused its inefficient genetic improvement and breeding. *E. nutans* harbors a pool of selected genes that enhance plant adaptation to cold, drought, and ultraviolet (UV) radiation (Liu et al., 2022). However, the *E. nutans* genome sequence is not yet available owing to its allohexaploid nature (StStYYHH) and large size (Lu, 1993). An *E. nutans* reference genome would facilitate research on this species and its adaptation to the Tibetan Plateau as well as comparative analysis of plants in the Triticeae.

**Figure 1.**
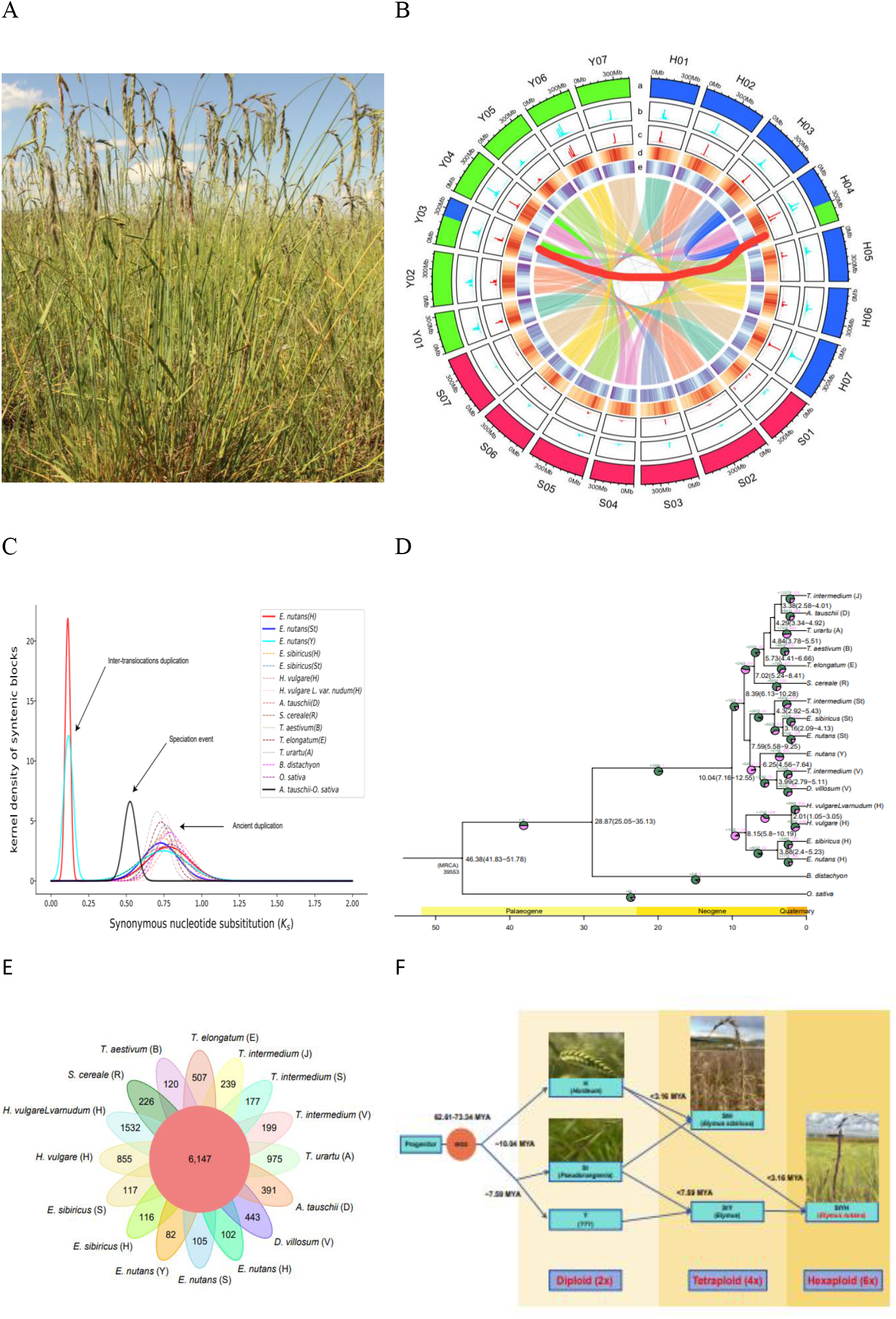
The reference genome of *Elymus nutans* provides insights into genome evolution and adaptation to the Qinghai-Tibetan Plateau. (A) *E. nutans* cv. Aba in the Tibetan Plateau. (B) Overview of the genomic features of the *E. nutans* genome. The outer to inner rows of the circle show (a) chromosome ideograms; (b) density of the LTR retrotransposons Cereba/Quinta; (c) Simple sequence repeat (SSR) density; (d) TE density; (e) high-confidence gene density. (C) Ks distribution for whole-genome duplication (WGD) events in *E. nutans* and other selected species. (D) Phylogenetic tree and divergence times of 12 related species. The branching time was estimated and shown based on the TimeTree data. The expansion (green) and contraction (red) gene families are also shown on the branches, with pie charts followed. (E) Venn diagram of gene families among subgenomes in Triticeae family members. (F) Speciation and evolutionary history of *E. nutans* with information about polyploidization and subgenome origins.

## Results and Discussion

### Genome assembly

Using advanced sequencing technology, we generated an allohexaploid reference genome for *E. nutans*, representing the three sets of chromosomes (subgenomes St, Y, and H). Initial contigs were assembled from long reads obtained using Oxford Nanopore Technology (ONT, 133.86×, N50> 29 kb; Supplemental Table 1), which were polished based on Illumina short reads (Supplemental Table 2). We assembled the contigs into 21 pseudo-chromosomes using Hi-C data (119.2×, Supplemental Table 3). After data cleaning and error correction, we obtained a final genome assembly of 9.36 Gb with a scaffold N50 of 424.5 Mb consisting of 21 chromosomes (Supplemental Table 4). The chromosomes were further grouped into three subgenomes (StStYYHH) based on similarity to the subgenomes of barley (*Hordeum vulgare*; HH) and *Elymus sibiricus* (StStHH) (Fig. 1B). The Benchmarking Universal Single-Copy Orthologs (BUSCO) score of the *E. nutans* assembly was 96.6%, and the LTR Assembly Index was 14.87–17.2, confirming its high quality. We successfully mapped 99.64% ONT and 97.1% NGS reads to the genome assembly, and the uniform coverage of mapped reads showed the relibility of the assembly, which was supported by the Hi-C heatmap. Synteny analysis revealed conservation among the three subgenomes, with one large reciprocal translocation detected between chromosomes H04 (175.1 Mb) and Y03 (153.8 Mb)(Fig. 1B). Collinearity between H04 and H03/St03 and between Y03 and Y04/St04 indicated the results from reciprocal translocation (Fig. 1B).

### Annotation of repetitive sequences

Among *E. nutans* genomic sequences, 83.89% were repetitive sequences (Supplemental Table 5), with up to 73.51% representing long terminal repeats (LTRs); the most abundant LTR categories were Copia and Gypsy. Gene annotation based on *de novo*, homology, and transcript-based predictions resulted in 114,214 gene models, including 39,341, 40,837, and 33,541 gene models for subgenomes H, St, and Y, respectively (Supplemental Table 4).We determined the locations of centromeric regions in the assembly based on enrichment of the LTR retrotransposons Cereba/Quinta, which are specific for centromeres in Triticeae genomes (Fig. 1B) and of centromeric SSRs in the maize (*Zea mays*) telomere-to-telomere genome (NCBI accession no. GCA_022117705.1) (Chen et al., 2023). The enrichment of transposable elements (TEs) matched the locations of centromeric regions and the centromeric and pericentromeric regions were gene-poor (Fig. 1B). We identified the centromeric regions on all 21 chromosomes, which will facilitate future studies of centromeres in Triticeae genomes.

### Related species evolution history

We explored the divergence of the three *E. nutans* subgenomes via sequence similarities with phylogenetically closely related species. The sequence identity in these species reached approximately 97.5% for subgenomes H and St (Supplemental Figure 1). Similar to other Gramineae species, the *Ks* values peaked at 0.7 to 0.82 (Fig. 1C), indicating that a whole-genome duplication (WGD) event affecting the three subgenomes that occurred approximately 62.61–73.34 million years ago (MYA). Another Ks peak at ∼0.1 was detected for subgenomes H and Y, likely due to the reciprocal translocation between H03 and Y04. From a phylogenetic tree reconstructed using 18 subgenomes of 12 species (Fig. 1D), we estimated the divergence time of the three subgenomes to be approximately 10.12 MYA, with H and St further splitting ∼7.49 MYA. Using divergence times and evolutionary relationships, we reconstructed a map for the speciation of *E. nutans*, estimating that hexaploid *E. nutans* (StStYYHH) was present <3.78 MYA (Fig. 1D) after the hybridization of an ancient diploid species (HH) and a tetraploid species (StStYY) (Fig. 1F; Liu et al., 2024). The history of the Y subgenome was traced to 6.25 MYA, when subgenome V in *Thinopyrum intermedium* and *Dasypyrum villosum* diverged from the ancestor of subgenomes Y and V (Fig. 1D). This finding differs from the recent conclusion that the origin of the Triticeae Y subgenome is from the St subgenome, based on chromosome-specific single-copy oligo probes (Chen et al., 2024).

### Comparative genomic analysis

Gene family analysis in *E. nutans* vs. nine other Triticeae subgenomes identified 102, 105, and 82 gene families unique to *E. nutans* subgenomes H, St, and Y, respectively, and 6147 gene families shared among subgenomes (Fig. 1E). More gene families expanded in subgenomes H (777 families) and St (756 families) than contracted (Fig. 1D), suggesting that gene expansion has played a key role in the adaptation of *E. nutans* to the Tibetan Plateau. These expanded gene families were significantly enriched in pathways related to environmental adaptation, with the conspicuous expansion of gene families associated with defense response processes in both H and St subgenomes (Supplemental Figure 2).

### Conclusion

In summary, our high-quality assembly of the three subgenomes of the Triticeae forage grass *E. nutans* provides critical insights into the evolutionary history of this species, which is uniquely adapted to extreme environmental conditions. This is the first study to our knowledge to confirm the origin of the Y subgenome, which shares a close relationship with the V subgenome. In addition to the large reciprocal translocation between chromosomes H04 and Y03 and the locations of the centromeres, we predict that *E. nutans* arose via two-step hybridization and that its speciation occurred ∼3.16 MYA.

## Materials and methods

### Genome sequencing

Fresh leaves were collected from an individual plant (*Elymus nutans* cv. Aba) at the 3-leaf stage at Sichuan Academy of Grassland Sciences, Hongyuan County, Sichuan Province, China (102.54°E, 32.79°N, 3495.86 m) and used for genome sequencing. Briefly, genomic DNA was extracted from the samples using a modified CTAB method (Agbagwa et al., 2012) and quantified using a Qubit 2.0 Fluorometer (Life Technologies, CA, USA) and an Agilent 2100 Bioanalyzer (Agilent Technologies, CA, USA). To generate short reads, two libraries were separately constructed with insert sizes of 270 bp and 8 kb using a TruSeq Nano DNA Library Prep Kit (Illumina, CA, USA) and an SQK-LSK108 Sequencing kit [Oxford Nanopore Technology (ONT), UK], respectively, and sequenced on an Illumina NovaSeq 6000 instrument. To generate long reads, the BluePippin™ System was first used to obtain large DNA segments of >30 kb, and a large-segment DNA library was prepared using an ONT Template prep kit (SQK-LSK109) and a NEB Next FFPE DNA Repair Mix kit (New England Biolabs, MA, USA). The high-quality library was sequenced on the ONT PromethION platform with Corresponding R9 cell and ONT Sequencing Reagents kit (EXP-FLP001.PRO.6). Hi-C libraries with insert sizes of 300–700 bp were constructed following a standard protocol described previously and sequenced on an Illumina HiSeq 4000 instrument.

Fresh tissues (leaf, stem, flower, seed, and root tissue) were collected from the same plant for total RNA extraction using a Plant total RNA Kit (TIANGEN, China). RNA-seq libraries were constructed using a TruSeq RNA Sample Preparation kit (Illumina) and sequenced on an Illumina HiSeq 4000 instrument. Raw sequencing data were processed by trimming adaptors and filtering out low-quality reads to generate clean data, which were used for annotation of protein-coding genes.

### Genome survey and assembly

A genome survey of *E. nutans* was performed using methods based on flow cytometry and *k*-mer frequency. *E. nutans* seeds were germinated in an incubator (germination conditions were set as follows: 25/20°C (day/night) under a 12-h light/12-h dark cycle, a light intensity of 225 ± 25 μmol m^−2^ s^−1^ and a relative humidity of 70% ± 5%), and root tips were excised from the seedlings and treated using the conventional pressing method (Yang et al., 2017) for karyotype analysis and chromosome counting. Genome size and ploidy were analyzed using a CyFlow Cube6 Flow Cytometer (Sysmex Partec, Germany) using *Triticum aestivum* „Chinese Spring” as a reference (Dolezel and Bartos, 2005). The Illumina short reads were used to estimate genome size, heterozygosity, and repeat content through *k*-mer frequency analysis (*k* = 23) using Jellyfish (Liu et al., 2013). The inferred size of the *E. nutans* genome is 10.50 Gb, with 0.01% heterozygosity and 78.00% repeat sequences.

*De novo* assembly of the *E. nutans* genome was based on ONT long reads. The ONT long reads were first corrected using Canu v1.7.1 with the parameter ErrorRate=0.025 (Koren et al., 2017), and the initial genome assembly was constructed using WTDBG v2.5 (Ruan and Li, 2020). The initial assembly was corrected three times using Recon (Vaser et al., 2017) based on the ONT long reads. The Illumina short reads were mapped to the initial assembly using the “MEM” module of BWA v0.7.10 (Li and Durbin, 2009), and three rounds of polishing were performed using Pilon with the parameters --mindepth 10 --changes --threads 4 --fix bases (Walker et al., 2014).

To construct pseudochromosomes, Hi-C raw data were trimmed using LACHESIS v2.0 (Burton et al., 2013). The high-quality Hi-C reads were then aligned to the draft assembly using BWA software, and uniquely mapped reads were selected by HiC-Pro v.2.10.0 (Servant et al., 2015) for further analysis. The manually corrected scaffolds were placed into 21 pseudochromosomes using LACHESIS software with the following parameters:

CLUSTER_MIN_RE_SITES=283, CLUSTER_MAX_LINK_DENSITY=2, ORDER_MIN_N_RES_IN_TRUNK=192, ORDER_MIN_N_RES_IN_SHREDS=178. Placement and orientation errors exhibiting obvious discrete chromatin interaction patterns were manually adjusted. The Hi-C read mapping and the quality of the final chromosome-level assembly were assessed using HiC-Pro software.

The completeness, of the final genome assembly was assessed using BUSCO v.5.0 (Simão et al., 2015) against the Embryophyta dataset with default parameters. Alignment and coverage of ONT long clean reads and Illumina short clean reads against the final genome assembly were evaluated using BWA software. LTR assembly index (LAI) was used to evaluate the continuity of the assembly based on full-length long terminal repeat retrotransposons (LTR-RTs) (Ou et al., 2018).

### Centromere and sub-genome identification

A reference-guided strategy based on the sequence collinearity of the subgenomes of *Hordeum vulgare* (HH) and *Elymus sibiricus* (StStHH) was used to distinguish subgenomes H, St, and Y of *E. nutans*. The 21 chromosomes were mapped to the *H. vulgare* genome using MUMmer v.3.0 (Kurtz et al., 2004) and clustered into seven groups, each containing three chromosomes. The seven chromosomes with the best alignments were selected as those from the H subgenome. Similarly, the 14 remaining chromosomes of *E. nutans* from subgenomes St and Y were mapped onto the H and St subgenomes of *E. sibiricus* (Yan et al., 2024) to identify the seven chromosomes from St, with the seven remaining chromosomes presumed to be from the Y subgenome. Overall, the *E. nutans* genome was resolved for the chromosomes and their numbers from subgenomes St, Y, and H.

The Cereba/Quinta sequences in *Triticum aestivum* were downloaded from NCBI (GenBank accession no. FN564437.1) to identify long terminal repeats in the *E. nutans* genome using RepeatMaster v.4.1.5 (http://www.repeatmasker.org/). The maize T2T genome assembly was downloaded (Chen et al., 2023), and the sequences in the centromere regions were retrieved based on published information. The maize centromeric sequences were aligned to the *E. nutans* genome, and the sequence hits were identified and summarized using a bin size of 1 Mb. The density distributions of the sequence hits were plotted onto the chromosomes, with the peaks used to determine the positions of the centromeres.

### Annotation of repetitive sequences and protein-coding genes

Repetitive sequences including tandem repeats and transposable elements (TEs) were identified from the whole genome of *E. nutans*. Tandem repeats were identified using GMATA v2.2 (Wang and Wang, 2016) and Tandem Repeats Finder v4.07b (Benson, 1999). A species-specific *de novo* repeat library was constructed using MITE-Hunter (Han and Wessler, 2010), LTR_FINDER v1.0.5 (Xu and Wang, 2007), and RepeatModeler v2.0.1 (https://github.com/Dfam-consortium/RepeatModeler). RepeatMasker software was then used to search for TEs against Repbase v19.06 (Jurka, 2005) and the species-specific *de novo* repeat library.

Protein-coding genes were predicted using an evidence-based annotation workflow by integrating evidence from *de novo* prediction, homology searches, and transcriptomic data. The *de novo* gene models were predicted using Augustus v.2.4 (Stanke et al., 2008) and SNAP software (Korf, 2004). For the homology-based approach, GeMoMa v.1.7 (Keilwagen et al., 2018) software was employed using reference gene models from other species, including *Arabidopsis thaliana, H. vulgare, Oryza sativa, Sorghum bicolor, T. aestivum*, and *Zea mays*. For transcript-based prediction, clean RNA-seq data were mapped to the *E. nutans* genome using HISAT v.2.0.4 (Kim et al., 2015) and assembled using StringTie v.1.2.3 (Pertea et al., 2015). GeneMarkS-T v.5.1 (Tang et al., 2015) was used to predict genes based on the assembled transcripts. PASA v.2.0.2 (Haas et al., 2008) was used to predict genes based on the unigenes assembled by Trinity v.2.11 (Grabherr et al., 2011). Gene models from these different approaches were combined using EVidenceModeler v.1.1.1 (Haas et al., 2008) and updated using PASA software after removing TE-related genes, pseudogenes, and noncoding genes using TransposonPSI v1.0 (Urasaki et al., 2017) with default settings.

The final gene models were annotated by searching the GenBank Non-Redundant (NR, data: 20200921), TrEMBL (data: 202005), Pfam v.33.1, SwissProt (data: 202005), eukaryotic orthologous groups (KOG, data: 20110125), and Kyoto Encyclopedia of Genes and Genomes (KEGG, data: 20191220) databases using BLAST software. Gene ontology (GO, data: 20200615) categories were annotated using Blast2go v.5.2.5 (Conesa et al., 2005) based on the NR annotation results.

### Comparative genomic analysis

The protein sequences of *E. nutans* (StStYYHH), *E. sibiricus* (StStHH), *H. vulgare* (HH), *Aegilops tauschii* (DD), *T. aestivum* B subgenome (BB), *Thinopyrum elongatum* (EE), *Secale cereale* (RR), *Triticum urartu* (AA), *H. vulgare* var. *nudum* (HH), *Brachypodium distachyon*, and *O. sativa* (as the outgroup) were aligned using Diamond v.0.9.29 (Buchfink et al., 2021) with an E-value of 0.001 using the all-vs-all strategy. The orthologous and paralogous gene families were identified by OrthoFinder v.2.4.0 (Emms and Kelly, 2019) and annotated using PANTHER v.14 (Mi et al., 2019). The protein sequences of single-copy orthologs were aligned using MAFFT v.7.205 (Nakamura et al., 2018), and a phylogenetic tree was reconstructed using IQ-TREE v.1.6.11 (Minh et al., 2020) based on the best evolution model JTT+F+I+G4 identified by ModelFinder (Kalyaanamoorthy et al., 2017) with 1000 bootstrap replicates. The divergence times were estimated using the MCMCtree program of PAML v.4.9 (Yang, 2007) with the parameters: burnin = 5000000, sampfreq=30, nsample=10000000. Expansion and contraction analysis of the gene families was performed using CAFE v4.2102 (Han et al., 2013). Three calibration points (*A. tauschii* vs. *O. sativa*: 42–52 MYA, *B. distachyon* vs. *A. tauschii*: 26–39 MYA, *T. aestivum* B subgenome vs. *T. urartu*: 2.6–5.3 MYA) were derived from the TimeTree database (http://www.timetree.org/).

Structural variations and syntenic relationship among the subgenomes of *E. nutans* and the related species *H. vulgare, T. elongatum, T. aestivum, T. urartu, A. tauschii*, and *O. sativa* were investigated using jcvi software (https://github.com/tanghaibao/jcvi/wiki). WGDI v. 0.6.5 (Sun et al., 2022) was used to identify the collinear blocks and to calculate the median Ks values for the blocks. The Ks curve for each species was fitted using WGDI software.

