## Supplemental Table 1 for "A reference genome assembly of the alpine forage grass *Elymus nutans*"

**Supplemental Table 1. Summary of Nanopore sequencing data**

| **Tissue** | **SeqNum** | **SumBase** | **N50Len** | **N90Len** | **MeanLen** | **MaxLen** | **MeanQual** |
| --- | --- | --- | --- | --- | --- | --- | --- |
| Leaf | 59,036,775 | 1,266,055,179,855 | 29,036 | 12,460 | 21,445 | 344,420 | 8.23 |

SeqNum: the number of sequencing reads SumBases: total bases of sequencing data; N50Len: N50 of sequencing data; N90Len: N90 of sequencing data; MeanLen: mean length of sequencing data; MaxLen: max length of sequencing data; MeanQual: mean quality of sequencing data.
