## Supplemental Table 2 for "A reference genome assembly of the alpine forage grass *Elymus nutans*"

**Supplemental Table 2. Summary of Illumina sequencing data**

| **Tissue** | **Library** | **ReadSum** | **BaseSum** | **GC (%)** | **Q30 (%)** |
| --- | --- | --- | --- | --- | --- |
| Stem/Leaf/Root | T01 | 96,294,374 | 28,846,649,322 | 57.12 | 95.08 |

ReadSum: total read number of sequencing data; BaseSum: total bases of sequencing data; GC(%): GC content; Q30(%): percentage of bases with Phred quality score ≥ 30.
