## Supplemental Table 3 for "A reference genome assembly of the alpine forage grass *Elymus nutans*"

**Supplemental Table 3. Summary of Hi-C sequencing data**

| **Tissue** | **Library** | **ReadSum** | **BaseSum** | **GC (%)** | **Q30 (%)** |
| --- | --- | --- | --- | --- | --- |
| Stem/Leaf/Root | H01 | 507,045,904 | 151,551,840,872 | 51.59 | 92.98 |
| Stem/Leaf/Root | H02 | 508,558,714 | 151,803,572,936 | 49.65 | 92.45 |
| Stem/Leaf/Root | H03 | 517,750,260 | 154,561,581,828 | 49.95 | 92.81 |
| Stem/Leaf/Root | H04 | 462,819,843 | 138,023,841,758 | 49.81 | 93.95 |
| Stem/Leaf/Root | H05 | 457,810,106 | 136,552,117,232 | 49.82 | 93.30 |
| Stem/Leaf/Root | H06 | 466,212,482 | 139,170,266,710 | 49.68 | 93.42 |
| Stem/Leaf/Root | H07 | 621,009,962 | 185,159,884,998 | 49.68 | 92.90 |

ReadSum: total read number of sequencing data; BaseSum: total bases of sequencing data; GC(%): GC content; Q30(%): percentage of bases with Phred quality score ≥ 30.
