## Supplemental Table 4 for "A reference genome assembly of the alpine forage grass *Elymus nutans*"

**Supplementary Table 4. Statistics for the genome assembly of *Elymus nutans***

|  | ***E. nutans**** | ***E. nutans H*** | ***E. nutans St*** | ***E. nutans Y*** | ***Sorghum bicolor*** | ***Triticum aestivum*** | ***Zea mays*** | ***Arabidopsis thaliana*** | ***Hordeum vulgare*** | ***Oryza sativa*** |
| --- | --- | --- | --- | --- | --- | --- | --- | --- | --- | --- |
| **GeneNum** | 114,214 | 39,341 | 40,837 | 33,541 | 34,129 | 107,544 | 35,615 | 27,381 | 39,718 | 38,852 |
| **GeneLen** | 390,341,448 | 133,468,086 | 141,411,111 | 114,370,622 | 126,726,290 | 375,144,978 | 157,589,684 | 60,368,916 | 238,572,893 | 110,901,512 |
| **AveGenlen** | 3,417.63 | 3,392.60 | 3,462.82 | 3,409.88 | 3,713.16 | 3,488.29 | 4,424.81 | 2,204.77 | 6,006.67 | 2,854.46 |
| **ExonLen** | 165,415,624 | 57,305,363 | 59,238,974 | 48,325,483 | 62,148,246 | 168,906,626 | 58,505,555 | 40,579,948 | 40,906,926 | 41,259,390 |
| **AveExonLen** | 1,448.30 | 1,456.63 | 1,450.62 | 1,440.79 | 1,820.98 | 1,570.58 | 1,642.72 | 1,482.05 | 1,029.93 | 1,061.96 |
| **ExonNum** | 528,136 | 180,165 | 187,495 | 158,794 | 166,869 | 502,038 | 170,559 | 145,401 | 154,975 | 160,012 |
| **CDSLen** | 145,823,238 | 50,520,939 | 52,161,198 | 42,647,313 | 39,640,380 | 133,312,040 | 41,331,000 | 33,343,421 | 40,906,926 | 41,259,390 |
| **AveCDSlen** | 1,276.75 | 1,284.18 | 1,277.30 | 1,271.50 | 1,161.49 | 1,239.60 | 1,160.49 | 1,217.76 | 1,029.93 | 1,061.96 |
| **CDSNum** | 514,468 | 175,478 | 182,131 | 155,193 | 153,976 | 477,350 | 160,069 | 140,295 | 154,975 | 160,012 |
| **IntronLen** | 224,925,824 | 76,162,723 | 82,172,137 | 66,045,139 | 64,578,044 | 206,238,352 | 99,084,129 | 19,788,968 | 197,665,967 | 69,642,122 |
| **AveIntronLen** | 1,969.34 | 1,935.96 | 2,012.20 | 1,969.09 | 1,892.18 | 1,917.71 | 2,782.09 | 722.73 | 4,976.74 | 1,792.50 |
| **IntronNum** | 413,922 | 140,824 | 146,658 | 125,253 | 132,740 | 394,494 | 134,944 | 118,020 | 115,257 | 121,160 |

GeneNum: number of genes; GeneLen: total gene length; AveGenelen: average gene length; ExonLen: total exon length; AveExonLen average exon length per gene; ExonNum: number of exons; CDSLen: total coding sequence (CDS) length; AveCDSlen: average CDS length per gene; CDSNum: number of CDSs; IntronLen: total intron length; AveIntronLen: average intron length per gene; IntronNum: number of introns. *The genes in *E. nutans* include the ones on the three subgenomes and unplaced contigs.
