## Supplemental Table 5 for "A reference genome assembly of the alpine forage grass *Elymus nutans*"

**Supplementary Table 5. Repetitive sequences in *Elymus nutans***

| **Repeat Class Description** | | | ***Elymus nutans*** | | ***Elymus nutans* H subgenome** | | ***Elymus nutans* St subgenome** | | ***Elymus nutans* Y subgenome** | |
| --- | --- | --- | --- | --- | --- | --- | --- | --- | --- | --- |
| **Repeat class** | **Type** | **Superfamily** | **bp** | **%** | **bp** | **%** | **bp** | **%** | **bp** | **%** |
| **Class I TEs** | **LTR** | **Copia** | 1,303,900,955 | 13.79 | 520,368,864 | 15.91 | 359,387,609 | 11 | 409,925,230 | 14.51 |
|  |  | **Gypsy** | 3,937,238,995 | 41.63 | 1,350,333,150 | 41.3 | 1,390,746,252 | 42.57 | 1,173,640,216 | 41.54 |
|  |  | **Unknown** | 591,340,798 | 6.25 | 241,523,820 | 7.39 | 213,458,654 | 6.53 | 131,609,961 | 4.66 |
|  | **nonLTR** | **LINE** | 31,231,910 | 0.33 | 13,459,742 | 0.41 | 9,229,518 | 0.28 | 8,421,629 | 0.3 |
|  |  | **SINE** | 813,401 | 0.01 | 119,480 | 0 | 349,688 | 0.01 | 341,246 | 0.01 |
| **Class II TEs** | **DNA_transposon** |  | 2,110,860 | 0.02 | 551,925 | 0.02 | 787,995 | 0.02 | 769,138 | 0.03 |
|  | **TIR** | **CACTA** | 1,209,659,372 | 12.79 | 324,248,132 | 9.92 | 441,425,583 | 13.51 | 425,370,393 | 15.05 |
|  |  | **Mutator** | 165,246,819 | 1.75 | 58,513,047 | 1.79 | 59,835,976 | 1.83 | 44,871,659 | 1.59 |
|  |  | **PIF_Harbinger** | 106,833,082 | 1.13 | 40,352,727 | 1.23 | 35,227,984 | 1.08 | 30,899,207 | 1.09 |
|  |  | **Tc1_Mariner** | 141,894,691 | 1.5 | 46,524,040 | 1.42 | 52,785,247 | 1.62 | 42,053,305 | 1.49 |
|  |  | **hAT** | 37,967,324 | 0.4 | 13,403,624 | 0.41 | 13,996,666 | 0.43 | 10,404,995 | 0.37 |
|  |  | **Unknown** | 12,359,774 | 0.13 | 4,090,763 | 0.13 | 4,247,328 | 0.13 | 3,986,806 | 0.14 |
|  | **nonTIR** | **Helitron** | 303,451,477 | 3.21 | 105,843,646 | 3.24 | 113,067,128 | 3.46 | 83,268,857 | 2.95 |
| **Repeat regions** |  |  | 90,348,161 | 0.96 | 27,216,380 | 0.83 | 34,066,244 | 1.04 | 25,574,199 | 0.91 |
| **Total** |  |  | 7,934,397,619 | 83.89 | 2,746,549,340 | 84 | 2,728,611,872 | 83.52 | 2,391,136,841 | 84.63 |
