## Supplemental Figure 1 for "A reference genome assembly of the alpine forage grass *Elymus nutans*"

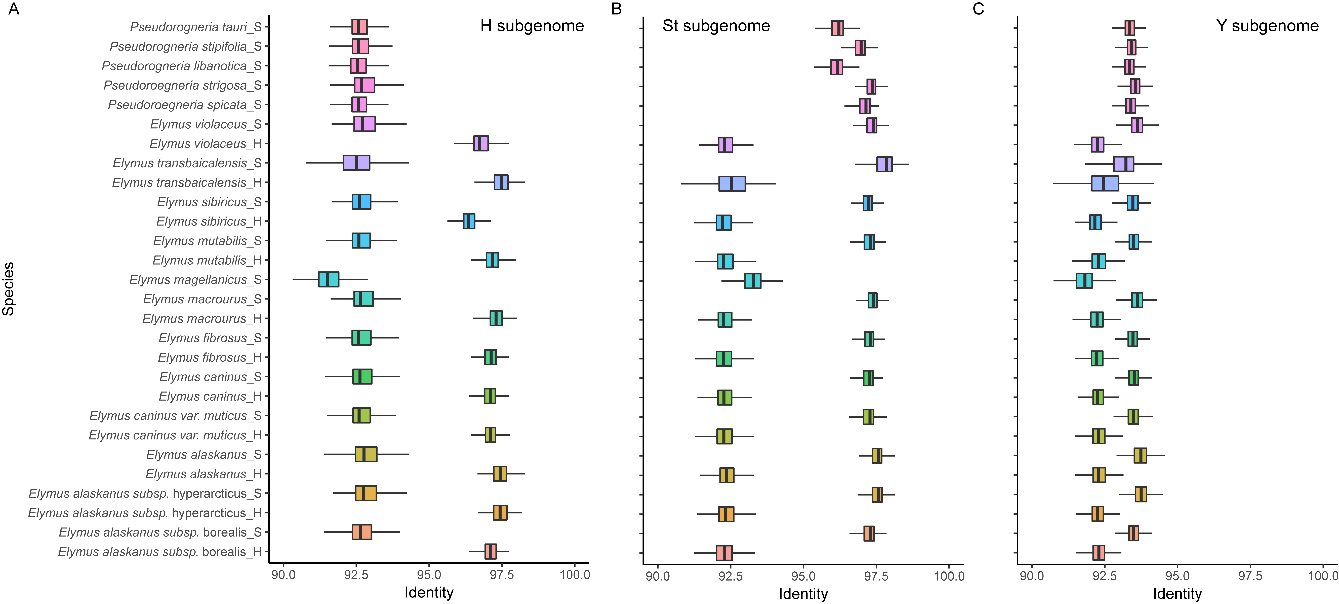

**Supplemental Figure 1. Sequence similarities of reads from different *Elymus* species that were uniquely mapped to the H (A), St (B), and Y (C) subgenomes of** ***E. nutans*.**
