## Supplemental Figure 2 for "A reference genome assembly of the alpine forage grass *Elymus nutans*"

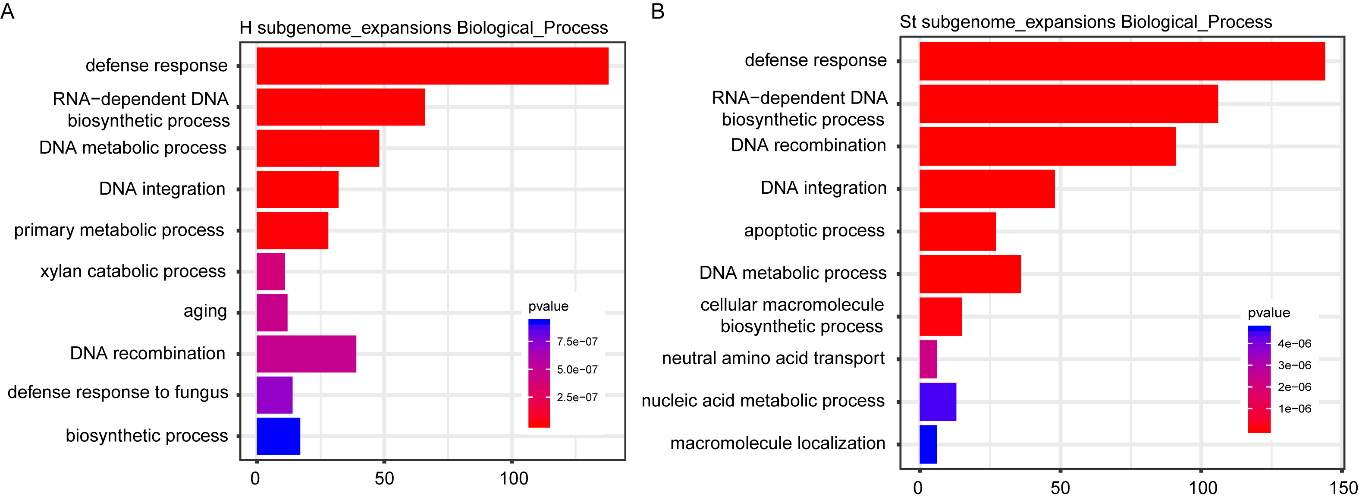

**Supplemental Figure 2. Top 10 most significantly enriched GO biological process terms for the expanded gene families in the H (A) and St (B) subgenomes.**
